# Assessing the Antimicrobial potentials of selected Plant Extracts against Post Harvest White Yam Rot

**DOI:** 10.1101/2024.09.05.611459

**Authors:** Aniefon A Ibuot, Iniobong I James

## Abstract

Antimicrobial potentials of selected aqueous and ethanolic plant extracts against post-harvest white yam rot was studied. Different concentrations (0.05%, 0.1%, 0.5%, 1.0%, 5.0% and 10%) of the selected plant extract *Azadirachta indica, Chromolaena odorata, Cymbopogan citratus, Ocimum gratissimum, Zingiber officinale* and *Allium sativum* were prepared. Fungi associated with post-harvest yam rot isolated and their percentage occurrence were; *F oxysporum (*23.7%), *Penicillium oxalicum* (27.55%), *Botryodiploidia theobromae* (10.98%), *Aspergillus niger* (17.89%), *Sclerotium rolfsii* (19.82%). The effect of selected plant extract on the growth of fungal isolates revealed ethanolic extract of *Allium sativum* having a very high growth inhibitory effect on all the fungal isolates which ranged from 71.8±0.98% to 100±0.00% at between 1.0% and 10% concentration of the plants extract. The aqueous plant extract of *Allium sativum* showed high inhibitory effect on all the fungal isolates but slightly lower than the ethanolic extract. Other plants extracts varied in their fungal inhibitory effect depending on the fungi. The phytochemical analysis of selected plants extracts showed the presence of most phytochemicals except for very few exceptions.

## 1. Introduction

White yam (*Dioscorea rotundata*) belongs to the family of *Dioscoreacea* and is monocotyledonous. It is among the most rated and easily accessed food crops of the tropical world. The varieties of yam that are edible are very important food crop and is a major carbohydrate staple food for millions of people leaving in West Africa. White yam is also a common food to people in Malaysia, Caribbean, some parts of Asia including parts of China and Japan (1). About three quarter of the total output of yams in the world is produced in Nigeria producing about 38.92 million metric tones annually (2).

Losses of yam tubers due to post harvest rot greatly affect the income of farmers and traders, threatens food security and also cause shortage of seed yams stored for planting. Yams losses in storage mostly due to rot tends to be heavy in Nigeria consequently, the demand for yam tubers has always exceeded its supply due to shortages. Between 20 and 39.5% of stored yam tubers may well be lost to rot microorganisms (3). In addition, an estimated 40% postharvest losses due to microbial attack in yam is reported

Some major microorganisms that causes yam rot in Nigeria are; *Fusarium solani, Rhizopus nodosus, Aspergillus niger, Clasdosporium pearospermum Fusarium monoliform*. Synthetic chemicals such as bleach, borax, sodium orthiophenyl phenate, captan, have been noted to reduce yam storage rot significantly. Regrettably because of the high cost of this chemicals farmers in most developing countries like Nigeria are unable to afford these synthetic chemicals for use. In addition, these synthetic chemicals accumulates in the ecosystem causing pesticide resistance to pathogens and also contaminates the water bodies causing huge health complications to the people (3).

The search for an eco-friendly, cheap botanical approach to preventing post-harvest yam rot therefore led to research on plants as potential tools for control of post-harvest yam rot.

## 2. Materials and Methods

### 2.1 Sample Collection and Preparation

Tubers of white yam showing various degrees of symptoms of dry rots were obtained from yam farmers from storage barns in Akwa Ibom State. The rotten yam were packed into polyethylene bag and taken to the laboratory for isolation and identification of fungal pathogens.

### 2.2 Isolation and identification of fungal pathogens

The infected/ rotten yam tubers were surface sterilized by dipping completely in a concentration of 5% sodium hypochlorite solution for 2min; the sterilized sections to be inoculated were then removed and rinsed in four successive changes of sterile distilled water (SDW).

Small sizes of approximately 2×2 mm was cut out with sterile scalpel from yam tubers infected with rot at inter-phase between the healthy and rotten portions of the tubers. The yam pieces will be placed on sterile filter paper in laminar Air flow cabinet to dry for 2 minutes.

A portion of the rotten yam was taken for serial dilution and there after cultivated in potato dextrose agar (PDA).

Sub-culturing of mycelia colonies from the inoculated plates was done.

Sterilized surgical blade was used to cut a section of mycelia and transferring the cut sections onto sterile PDA plates. The plates (inoculated) were incubated at ambient room temperature (30±5^0^C) for 2-3 days. The purified isolates kept in slants and stored for characterization and pathogenicity test. Microscopic examination, morphological characteristics and identification was done (4-5)

### 2.3 Pathogenicity test of the isolated fungi

Pathogenicity test of the isolated fungi was carried out using the method of Okigbo and Ikediugwu, (1) Pathogenicity test of the four test fungal isolates *F oxysporum, Penicillium oxalicum, Botryodiploidia theobromae, Aspergillus niger, Sclerotium rolfsii* from rotten yam tubers were tested on their ability to or not to cause rot on healthy yam tubers. Fresh healthy white yam tubers were washed with distilled water and sterilized using 70% ethanol. Aseptically, a cylindrical 1cm deep was cut out from the healthy white yam tuber using a sterile 5mm cork borer. A sample of the isolate from a 5 day old PDA pure culture was inserted into the hole created and was sealed. The replaced part at the point of inoculation was covered with sterile petroleum jelly; the process was repeated for the control without any pathogen. The inoculated white yam tubers were incubated at room temperature for a period of 7-14 days; signs of rot were observed daily. After 14days of incubation, the yam tubers were cut open through the inoculation point of each to reveal the inner portion and the level of rot was determined.

### 2.4 Preparation of plants extracts

The method of Taiga *et al*., (6) was used with some modifications in extraction and preparation of extract of *Azadirachta indica, Chromolaena odorata, Cymbopogan citratus, Ocimum gratissimum, Zingiber officinale, Allium sativum*

### 2.4 Phytochemical Screening of the plants

Test for the presence of alkaloids, flavonoids, steroid, tannin, phenol, saponin, cardiac glycosides, terpenes and steroids, resins was carried out using the method of Edeoga et al., (7) and Oluduro, (8)

### 2.5 Effect of plant extracts on fungal pathogens

Direct medium treatment was done using the method of Gwa and Ekefan (9); Gwa and Akombo, (10). The effect of selected plant extracts on fungal growth was examined using the growth inhibition test invitro. One ml of each of the plant extract concentrations was dispensed into 9ml of melted PDA in a Petri dish, gently shaken and allowed to solidity; this was used in preparing 0.05%, 0.1%, 0.5%, 0.1%, 1%, 5% and 10% plants extracts concentrations. A portion from the colony of 5 -day old pure cultures (mycelia disc) of each of the test fungal isolates was inoculated onto the center of the PDA-plant extract mixture plate. Negative control experiments were carried out with no addition of any plant extract and the positive control PDA plates had 1 ml of 0.5 g Mancozeb added to it. This was done in triplicates and the plates incubated at 28^0^C for five days. Percentage inhibition of fungal growth was calculated using the formula below:

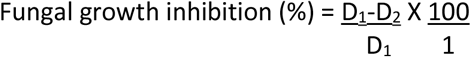

Where D_1_ – Average Diameter of control and D_2_ – Average Diameter of fungal colony with treatment

### 2.6 Statistical Analysis

Complete Randomized Design was used as the experimental design with samples in triplicates. Analysis of variance was carried out and Duncan letters at 0.05 probability level indicating significant levels of values was used in separating means

## 3.0 Result

Fungi frequently associated with yam tuber rot in this study were *F oxysporum Penicillium oxalicum, Botryodiploidia theobromae, Aspergillus niger* and *Sclerotium rolfsii with* F. oxysporum as the most prevalent organism (Table 1)

**Table 1.** Percentage (%) occurrence of fungi in rotted yam.

| Isolates | Occurrence (%) |
| --- | --- |
| *F oxysporum* | 23.76 |
| *Penicillium oxalicum* | 27.55 |
| *Botryodiploidia theobromae* | 10.98 |
| *Aspergillus niger* | 17.89 |
| *Sclerotium rolfsii* | 19. |

The ethanolic extract of *A. sativum* at 1%, 5% and 10% concentrations had 100±0.00% fungal growth inhibition on *Botryodiplodia theobromae*. Ethanolic plant extract of *A. indica* at 1%, 5% and 10% concentration had 79.8±2.19% to 100±0.00%. Ethanolic plant extract of *C. citratus*.at 1%, 5% and 10% concentrations showed a fungal growth inhibition that ranged from 80.9±6.11% to 96±2.57% (Table 7). The ethanolic extract of *C. odorata* and *O. gratissimum* showed very minimal fungal growth inhibition which ranged between 7.1±3.23 % to 14.7±5.12% (Table 7). The aqueous plant extract of these plants revealed slight reduction in the fungal growth inhibition compared to the ethanolic extract (Table 2).

**Table 2.** Effect of aqueous plant extracts on the fungal growth inhibition of *Botryodiplodia theobromae*incubated at 28^0^C for 5 days.

|  | Fungal growth inhibition (%) at specific extract concentration |  |  |  |  |  |
| --- | --- | --- | --- | --- | --- | --- |
|  | 0.05% | 0.1% | 0.5% | 1% | 5% | 10% |
| Allium sativum | 9.2±0.78 | 28.3±4.00 | 88.5±11.00 | 100±0.00 | 100±0.00 | 100±0.00 |
| Azadirachta indica | 3.4±1.98 | 8.0±3.87 | 26.5±1.20 | 70.6±7.90 | 76.4±6.09 | 80.2±4.89 |
| Chromolaena odorata | 0.0±0.00 | 0.0±0.00 | 0.0±0.00 | 1.9±3.89 | 1.9±4.99 | 2.1±6.09 |
| Cymbopogan citratus | 4.3±2.89 | 10.0±1.90 | 19.9±1.30 | 60.2±6.90 | 65.6±5.78 | 65.9±2.98 |
| Ocimum gratissimum | 0.0±0.00 | 0.0±0.00 | 0.8±2.89 | 2.7±4.98 | 2.9±6.00 | 3.6±5.10 |
| Zingiber officinale | 0.0±4.00 | 0.0±0.00 | 8.2±3.10 | 39.±9.10 | 39.6±1.89 | 39.7±4.19 |
| + ve control | 80.0±1.87 | 83.0±2.19 | NA | NA | NA | NA |
| -ve control | 0.00±0.00 | 0.00±0.00 | 0.00±0.00 | 0.00±0.00 | 0.00±0.00 | 0.00±0.00 |

*Results are in Mean ±Std*

The ethanolic plant extract of *A. sativum* at 1%, 5% and 10% concentrations had 100±0.00 % fungal growth inhibition on *Penicillium oxalicum*. Ethanolic plant extract of *A*.*indica* at 1%, 5% and 10% concentrations had a range of 70.2± 5.22% to 100±0.00% fungal growth inhibition and ethanolic plant extract of *O. gratissimum* at 1%-10% concentration had fungal growth inhibition which ranged from 70.1±3.19% to 90.1±3.17%. The ethanolic plant extract of *O. odorata, C. citratus* and *Z. officinale* at 5% to 10% concentrations had fungal growth inhibition which ranged from 61.0±1.22% to 78.0±2.78% (Table 8) while the aqueous plant extract of these plants extracts showed lower fungal growth inhibitory effect (Table 3).

**Table 3.** Effect of aqueous plant extracts on the growth inhibition of Penicillium oxalicum incubated at 28^0^C for 5 days.

|  | Growth inhibition (%) at specific extract concentration |  |  |  |  |  |
| --- | --- | --- | --- | --- | --- | --- |
|  | 0.05% | 0.1% | 0.5% | 1% | 5% | 10% |
| Allium sativum | 9.4±6.09 | 20.0±6.01 | 75.1±0.49 | 100±0.00 | 100±0.00 | 100±0.00 |
| Azadirachta indica | 0.0±2.19 | 2.0±3.10 | 24.4±7.10 | 57.5±4.10 | 59.2±7.01 | 61.2±6.80 |
| Chromolaena odorata | 0.0±0.00 | 0.0±0.00 | 20.3±4.87 | 38.0±0.99 | 41.0±2.46 | 42.0±7.49 |
| Cymbopogan citratus | 0.0±0.00 | 0.0± 0.00 | 20.0±7.86 | 46.9±2.10 | 48.0±6.82 | 48.0±0.19 |
| Ocimum gratissimum | 9.5±3.10 | 24.8±5.02 | 31.0±1.80 | 58.8±4.00 | 61.9±8.01 | 71.5±6.76 |
| Zingiber officinale | 0.0±0.00 | 0.7±1.83 | 6.0±5.80 | 40.5±1.98 | 42.8±2.67 | 46.6±4.20 |
| +ve control | 81.0±0.93 | 82.00±0.98 | NA | NA | NA | NA |
| -ve control | 0.00±0.00 | 0.00±0.00 | 0.00±0.00 | 0.00±0.00 | 0.00±0.00 | 0.00±0.00 |

*Results are in Mean ±Std*

**Table 4.** Effect of aqueous plant extracts on the growth inhibition of *Fusarium oxysporum* incubated at 28^0^C for 5 days.

|  | Growth inhibition (%) at specific extract concentration |  |  |  |  |  |
| --- | --- | --- | --- | --- | --- | --- |
|  | 0.05% | 0.1% | 0.5% | 1% | 5% | 10% |
| *Allium sativum* | 10.2±2.10 | 38.8±0.98 | 50.1±0.98 | 71.8±0.98 | 100±0.00 | 100±0.00 |
| *Azadirachta indica* | 9.2±5.09 | 21.4±2.10 | 32.0±2.10 | 62.8±3.09 | 71.0±0.98 | 76.5±0.89 |
| *Chromolaena odorata* | 1.4±9.01 | 3.9±1.87 | 25.0±6.01 | 30.5±7.05 | 31.2±7.76 | 36.4±2.10 |
| *Cymbopogan citratus* | 6.2±4.01 | 15.1±4.98 | 25.0±3.19 | 61.5±8.09 | 64.5±7.01 | 68.0±6.01 |
| *Ocimum gratissimum* | 1.4±0.89 | 4.3±2.09 | 14.2± 7.01 | 43.0±3.09 | 46.0±2.09 | 49.1±7.93 |
| *Zingiber officinale* | 7.9±1.09 | 23.7±5.10 | 50.2±2.09 | 62.0±2.09 | 66.0±1.09 | 71.0±8.01 |
| +ve control | 80.7±2.10 | 80.7±7.90 | NA | NA | NA | NA |
| -ve control | 0.00±0.00 | 0.00±0.00 | 0.00±0.00 | 0.00±0.00 | 0.00±0.00 | 0.00±0.00 |

*Results are in Mean ±Std*

**Table 5.**
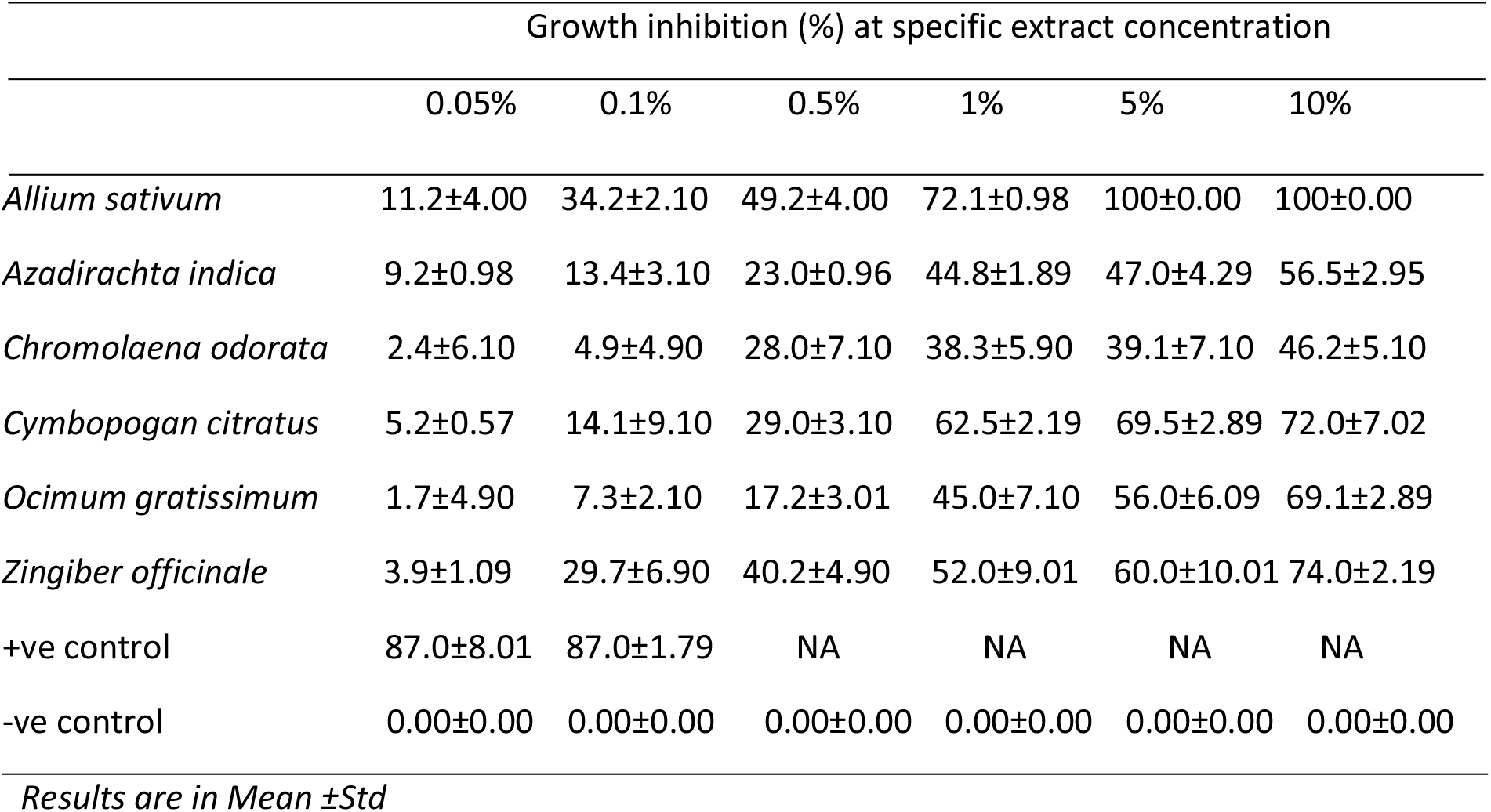
Effect of aqueous plant extracts on the growth inhibition of *Aspergiluus niger* incubated at 28^0^C for 5 days.

|  | Growth inhibition (%) at specific extract concentration |  |  |  |  |  |
| --- | --- | --- | --- | --- | --- | --- |
|  | 0.05% | 0.1% | 0.5% | 1% | 5% | 10% |
| *Allium sativum* | 11.2±4.00 | 34.2±2.10 | 49.2±4.00 | 72.1±0.98 | 100±0.00 | 100±0.00 |
| *Azadirachta indica* | 9.2±0.98 | 13.4±3.10 | 23.0±0.96 | 44.8±1.89 | 47.0±4.29 | 56.5±2.95 |
| *Chromolaena odorata* | 2.4±6.10 | 4.9±4.90 | 28.0±7.10 | 38.3±5.90 | 39.1±7.10 | 46.2±5.10 |
| *Cymbopogan citratus* | 5.2±0.57 | 14.1±9.10 | 29.0±3.10 | 62.5±2.19 | 69.5±2.89 | 72.0±7.02 |
| *Ocimum gratissimum* | 1.7±4.90 | 7.3±2.10 | 17.2±3.01 | 45.0±7.10 | 56.0±6.09 | 69.1±2.89 |
| *Zingiber officinale* | 3.9±1.09 | 29.7±6.90 | 40.2±4.90 | 52.0±9.01 | 60.0±10.01 | 74.0±2.19 |
| +ve control | 87.0±8.01 | 87.0±1.79 | NA | NA | NA | NA |
| -ve control | 0.00±0.00 | 0.00±0.00 | 0.00±0.00 | 0.00±0.00 | 0.00±0.00 | 0.00±0.00 |

*Results are in Mean ±Std*

**Table 6.** Effect of aqueous plant extracts on the growth inhibition of *Sclerotium rolfsii* incubated at 28^0^C for 5 days.

|  | Growth inhibition (%) at specific extract concentration |  |  |  |  |  |
| --- | --- | --- | --- | --- | --- | --- |
|  | 0.05% | 0.1% | 0.5% | 1% | 5% | 10% |
| *Allium sativum* | 3.6±2.10 | 8.7±1.09 | 89.0±4.09 | 90.7±1.06 | 98.0±0.98 | 100.0±3.78 |
| *Azadirachta indica* | 9.5±2.98 | 27.3±4.09 | 33.2±5.89 | 69.7±4.19 | 79.1±7.09 | 89.4±2.10 |
| *Chromolaena odorata* | 0.00±4.28 | 1.7±5.90 | 2.8±0.79 | 8.7±3.78 | 9.8±5.90 | 13.7±2.98 |
| *Cymbopogan citratus* | 2.5±0.89 | 10.3±5.89 | 38.1±1.78 | 61.2±6.09 | 68.9±5.98 | 70.1±2.09 |
| *Ocimum gratissimum* | 1.4±3.98 | 5.1±2.19 | 9.3±0.99 | 10.0±4.67 | 12.1±2.09 | 13.3±2.08 |
| *Zingiber officinale* | 1.3±3.85 | 3.5±2.64 | 14.3±7.09 | 18.2±6.78 | 29.6±0.78 | 38.6± 0.98 |
| +ve control | 86.0±3.29 | 86.0±1.09 | NA | NA | NA | NA |
| -ve control | 0.00±0.00 | 0.00±0.00 | 0.00±0.00 | 0.00±0.00 | 0.00±0.00 | 0.00±0.00 |

*Results are in Mean ±Std*

**Table 7.** Effect of ethanolic plant extracts on the growth inhibition of *Botryodiplodia theobromae*incubated at 28^0^C for 5 days.

|  | Growth inhibition (%) at specific extract concentration |  |  |  |  |  |
| --- | --- | --- | --- | --- | --- | --- |
|  | 0.05% | 0.1% | 0.5% | 1% | 5% | 10% |
| *Allium sativum* | 13.3±1.24 | 38.0±3.45 | 89.8±5.56 | 100±0.00 | 100±0.00 | 100±0.00 |
| *Azadirachta indica* | 4.4±7.11 | 20.7±3.23 | 42.3 ±2.12 | 79.8 ±2.19 | 100±0.00 | 100±0.00 |
| *Chromolaena odorata* | 3.0±1.89 | 4.0±3.18 | 5.8± 2.78 | 7.5±2.58 | 8.1± 5.12 | 8.8±3.12 |
| *Cymbopogan citratus* | 5.2±4.12 | 19.5±41 | 31.4±4.11 | 80.9±6.11 | 88.0±5.10 | 96.4±2.57 |
| *Ocimum gratissimum* | 1.0±7.11 | 3.5±3.10 | 5.8±1.88 | 7.1±3.23 | 9.1±3.44 | 14.7±5.12 |
| *Zingiber officinale* | 0.5±3.18 | 2.1±2.10 | 29.1±2.30 | 52.0±2.19 | 59.1±3.52 | 68.8±1.22 |
| + ve control | 80.0±1.56 | 83.0±1.89 | NA | NA | NA | NA |
| -ve control | 0.00±0.00 | 0.00±0.00 | 0.00±0.00 | 0.00±0.00 | 0.00±0.00 | 0.00±0.00 |

*Results are in Mean ±Std*

**Table 8.** Effect of ethanolic plant extracts on the growth inhibition of *Penicillium oxalicum* incubated at 28^0^C for 5 day.

|  | Growth inhibition (%) at specific extract concentration |  |  |  |  |  |
| --- | --- | --- | --- | --- | --- | --- |
|  | 0.05% | 0.1% | 0.5% | 1% | 5% | 10% |
| Allium sativum | 10.3±4.33 | 38.2±2.11 | 86.4±1.98 | 100±0.00 | 100±0.00 | 100±0.00 |
| Azadirachta indica | 4.0±2.12 | 14.9±3.11 | 47.9±0.91 | 70.2±5.22 | 78.9±3.11 | 100±6.11 |
| Chromolaena odorata | 3.7±3.11 | 12.0±1.98 | 39.7±2.11 | 59.8±3.18 | 61.0±1.22 | 72.0±4.11 |
| Cymbopogan citratus | 3.0 ±0.98 | 10.5±2.18 | 35.9±4.11 | 62.5±2.11 | 70.1±3.12 | 78.8±3.92 |
| Ocimum gratissimum | 10.8±2.89 | 39.9±1.77 | 50.0±1.88 | 70.1±3.19 | 83.2±6.11 | 90.1±3.17 |
| Zingiber officinale | 2.0±0.89 | 8.1±4.00 | 30.0±2.56 | 48.2±9.12 | 59.0±3.18 | 70.0±2.78 |
| +ve control | 81.0±1.09 | 82.0±5.12 | NA | NA | NA | NA |
| -ve control | 0.00 ±0.00 | 0.00±0.00 | 0.00±0.00 | 0.00±0.00 | 0.00±0.00 | 0.00±0.00 |

*Results are in Mean ±Std*

The ethanolic plant extract of *A. sativum, A indica, C citratus* and *Z. officinale* at 1%, 5% and 10% concentrations had fungal growth inhibition on *Fusarium oxysporium* which ranged from 73.2±4.00% to 100±0.00%, with *A sativum* having the most impressive inhibitory effect (Table 9). Ethanolic plant extract of *C. odorata* and *O. gratissimum* at 1%, 5% and 10% concentrations had lower inhibitory effect which ranged from 40.2±7.11% to 62.9±4.16% (Table 9). The aqueous plant extract of the plants in their respective concentrations had lower fungal growth inhibition on *Fusarium oxysporium* compared to the ethanolic plants extracts (Table 4).

The ethanolic plant extracts of *A. sativum, C. citratus, O. gratissimum* and *Z. officinale* at 1%, 5% and 10% concentration had fungal growth inhibition on *A. niger* which ranged from 56.2±2.19% to 100±0.00%. *A. sativum* was observed to have the most impressive inhibitory effect (Table 10) while the aqueous plant extract of same plants had slightly less inhibitory effect on *A. niger* (Table 5). Ethanolic extract of *A. indica* and *C. odorata* at 1%. 5% and 10% concentrations had fungal growth inhibition which ranged from 43.6±7.11% to 65.1±8.00% (Table 10), while the inhibitory effect of the aqueous plant extract were observed to be slightly lower as compared to the ethanolic plant extract (Table 5)

**Table 9.**
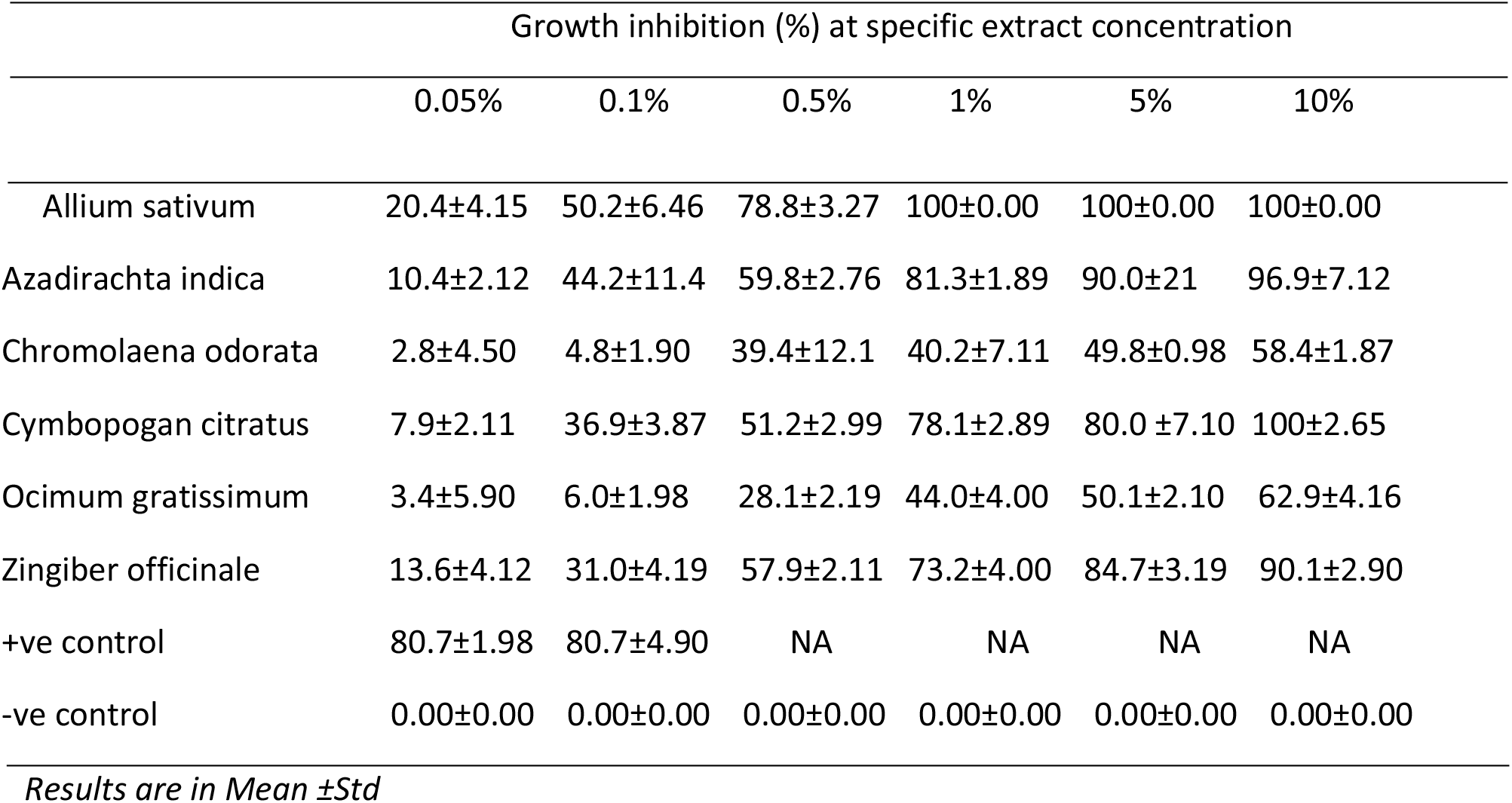
Effect of ethanolic plant extracts on the growth inhibition of *Fusarium oxysporum* incubated at 28^0^C for 5 days.

**Table 10.**
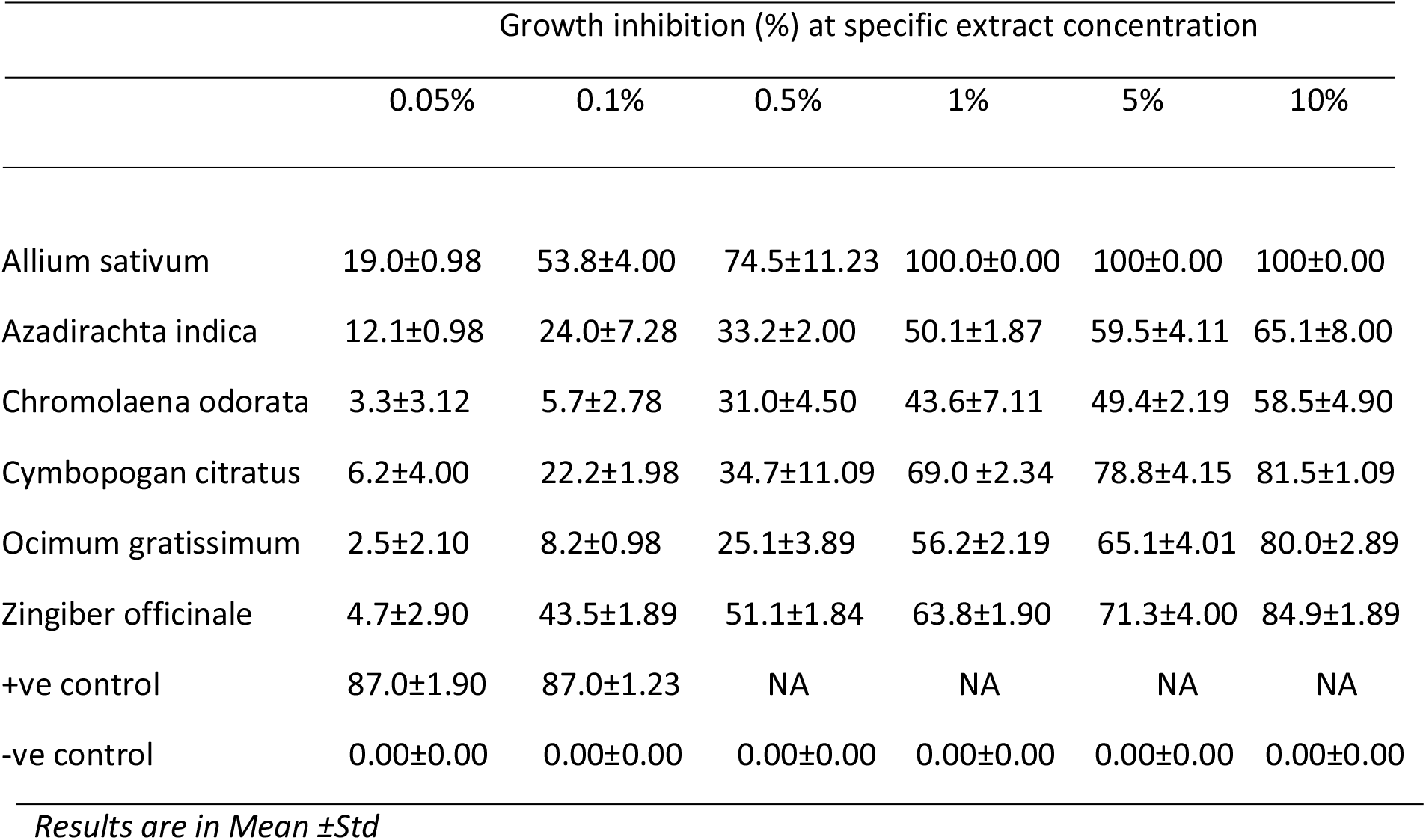
Effect of ethanolic plant extracts on the growth inhibition of *A. niger* incubated at 28^0^C for 5 days.

The ethanolic plant extracts of *A. sativum, A. indica* and *C. citratus* at 1%, 5% and 10% concentrations was observed to have a fungal growth inhibition on *S. rolfsii* which ranged from 70±1.29% to 100±0.00%, with

*A. sativum* showing a very impressive inhibitory effect at 0.05% concentration compared to other plants extracts (Table 11). The aqueous plant extracts of same plants showed slightly low inhibitory effect compared to the ethanolic counterparts (Table 6). Ethanolic plant extracts of *C. odorata, O. gratissimum* and *Z. officinale* at 1%, 5% and 10% concentration showed a fungal growth inhibition on *S. rolfsii* ranging from 9.1±4.11% to 42.5±1.98% (Table 11). The aqueous plant extract of the plants showed slightly lower inhibition on *S. rolfsii* compared with the ethanolic plant extracts (Table 6).

**Table 11.**
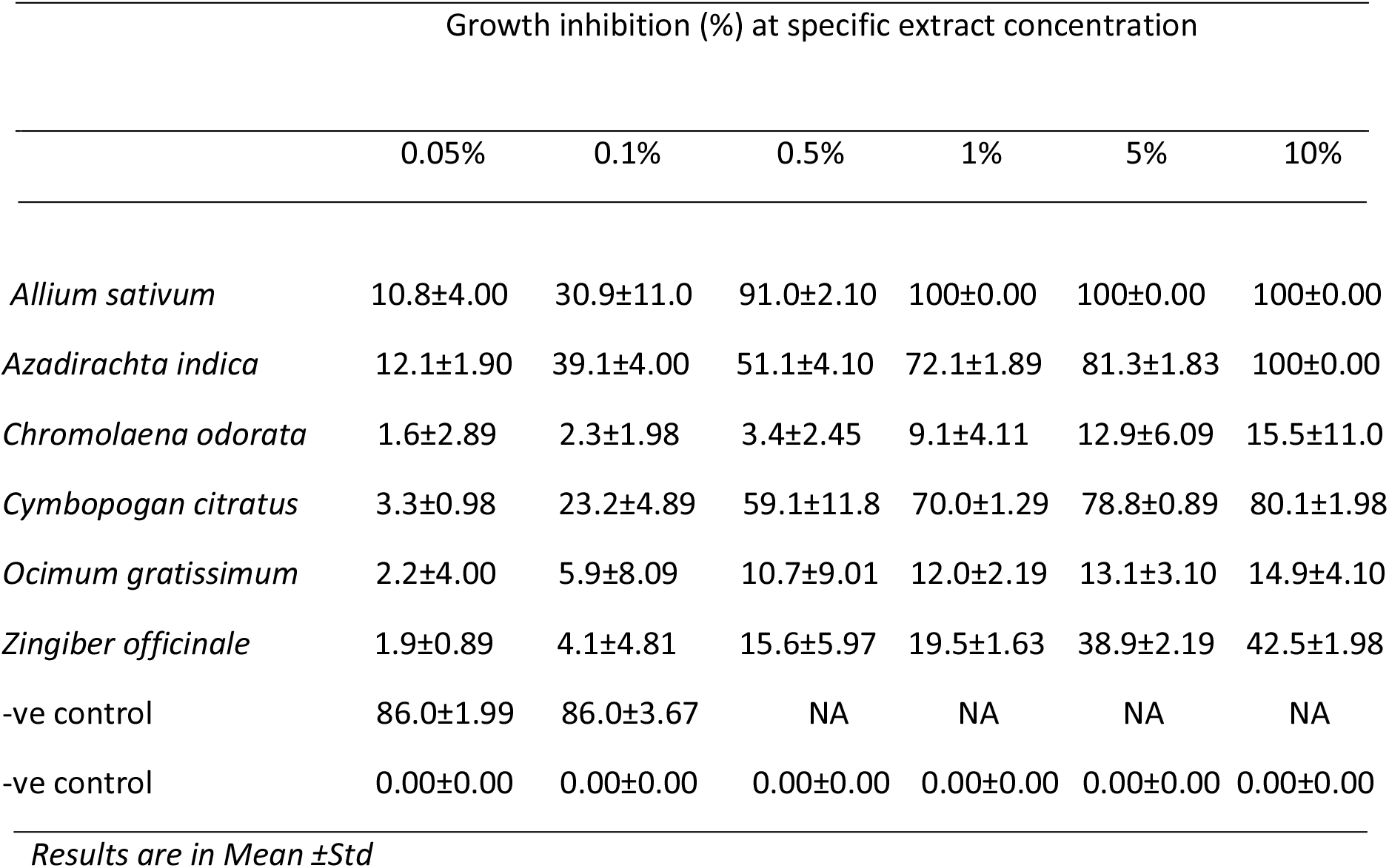
Effect of ethanolic plant extracts on the growth inhibition of *Sclerotium rolfsii* incubated at 28^0^C for 5 days.

The phytochemical screening of test plants shows that most of the phytochemicals tested are present in the test plants with few exceptions (Table 12).

**Table 12.** Phytochemical composition of test plants extract.

|  | *Allium sativum* | *Azadirachta indica* | *Chromolaena odorata* | *Cymbopogon citratus* | *Ocimum gratissimum* | *Zingiber officinale* |
| --- | --- | --- | --- | --- | --- | --- |
| Alkaloids | + | + | + | + | + | + |
| Flavonoids | + | + | + | + | + | + |
| Tannins | - | - |  | + | + | + |
| Saponins | - | + | + | + | + | + |
| Balsam | - | - |  | - | + | - |
| Cardiac glycosides | + | + | + | — | — | — |
| Terpenes and steroids | + | + | - | + | — | + |
| Resin | + | - | + | - | + | + |
| Phenols | - | + | + | + | - | + |

## 4.0 Discussion

Several studies have been carried out on post-harvest yam tuber rot caused by microorganisms from the farm to the barns where yams are stored (11-12) The microorganisms associated with post-harvest yam rot have been identified (11,1,13) which is in agreement with the ones identified in this work. Severally, these organisms are able to access yam tubers through natural openings and injuries that occurs in the process of harvesting and transporting from the farm to the barns where they are stored (11). Besides, yam tuber may have already been infected by pathogens from the diseases of leaves or roots. The soil also is a reservoir of microorganism, hence the soil particles clustering on the tuber can expose the tuber to pathogens of yam rot. Extracts from six plants were employed in these research to formulate cheap, eco-friendly and simpler botanical approach to control yam rot. This research reveals that the most impressive botanical fungicide extracts was extract of *Allium sativum* whose ethanolic extract showed 100% fungal growth inhibition against the all the fungi used in this study. *Azadiracta indica, Ocimum gratissimum, C. citritrus* and *Z. offinicale* also showed significant fungal growth inhibition. The result agrees with the reports of Okigbo and Ogbannaya (3), Okigbo and Nmeka (14). *Allium sativum* was observed to inhibit fungal growth of fungi used in this study at very minimum concentrations.

Shashiskanth et al., (15) reports that *Allium sativum* contains allicin as its main biological compound which is reported to be soluble in water and can inhibit the growth of broad range of pathogenic organisms.

Awuah (16) also reported that *Ocimum gratissimum* showed inhibitory properties in the control of yam rot organism which is evident in this work. The plant array used in this study is relatively available and cheap to prepared hence can be appreciated by small scale farmers. These extract are also known to be eco-friendly and non-toxic to man compared to the synthetic chemicals.

The phytochemical screening of test plants shows that most of the phytochemicals tested are present in the test plants with few exceptions. Biologically active components present in plants have been reported to bestow resistance to plants against pathogenic microorganisms (17). Hence the fungal growth inhibitory effect of these plants could be connected to the presence of various phytochemical component in the plants (18)

## 5.0 Conclusion

The test plant extracts *Allium sativum, Azadirachta indica, Chromolaena odorata, Cymbopogan citratus Ocimum gratissimum, Zingiber officinale* showed impressive inhibitory effect against pathogens of post-harvest white yam rot. These clearly reveals that these plants extracts are becoming a more reliable botanical tool and could be used to effectively prevent post-harvest yam rot and to increase the shelf life of white yam tubers. Therefore efforts should be made towards exploring the vast potential of these plant extracts for effective prevention of post-harvest yam rot to enhance food security.

## Acknowledgements

The article was sponsored by TETFund research grant, Nigeria

## Disclosure of conflict of interest

The authors declare no conflict of interest

## Notes

### Competing Interest Statement

The authors have declared no competing interest.

